# Muscle-Specific ECM Fibers Made with Anchored Cell Sheet Engineering Support Tissue Regeneration in Rat Models of Volumetric Muscle Loss

**DOI:** 10.1101/2024.12.15.628541

**Authors:** Alireza Shahin-Shamsabadi, John Cappuccitti

**Affiliations:** Evolved.Bio, 280 Joseph Street, Kitchener, Ontario, Canada

**Keywords:** Scaffold-free Biofabrication, Anchored cell sheet engineering, Volumetric muscle loss, Animal study, Quantitative image analysis

## Abstract

Volumetric muscle loss (VML) represents a critical unmet need in regenerative medicine, with no established standard of care. This study introduces a novel therapeutic strategy using tissue-specific skeletal muscle extracellular matrix (ECM) fibers fabricated using scaffold-free Anchored Cell Sheet Engineering technology. These engineered fibers replicate the native ECM composition and microarchitecture of skeletal muscle, incorporating essential structural and basement membrane proteins. In a rat VML model, engineered ECM fibers demonstrated a promising regenerative capacity compared to commercial porcine-derived small intestine submucosa (SIS) ECM. Over an 8-week period, the engineered fibers preserved muscle volume and weight, regulated inflammatory and fibrotic responses, and promoted vascularization. In contrast, SIS was rapidly degraded by week 4 and associated with excessive fibrotic response. Force recovery in the muscles treated with engineered ECM fibers was lower at the 8-week time point (77% compared to 91% in the control group), but histological and immunohistochemical analyses revealed newly formed, dispersed muscle fibers exclusively within the repaired muscle tissue treated with engineered ECM fibers. Importantly, only in cases where engineered ECM fibers were used, muscle weight was preserved, resulting in similar normalized force-to-weight recovery across all groups (87% in the test group vs. 88% in the control group). The histological analyses further demonstrated ongoing tissue remodeling, indicative of sustained regeneration, in contrast to the premature fibrotic healing observed in the other groups. A novel quantitative image analysis workflow using a custom Python script, enabled objective assessment of spatial tissue heterogeneity through histology and immunohistochemistry images, setting a new standard for tissue regeneration analysis. These findings establish engineered tissue-specific ECM fibers as a transformative approach for VML treatment and lay the groundwork for translation to clinical applications.

## 1. Introduction

Skeletal muscle accounts for approximately 40% of human body mass and is characterized by its highly organized structure and remarkable regenerative capacity. This regenerative potential is primarily driven by the activation of resident muscle stem cells, known as satellite cells, and their interactions with the extracellular matrix (ECM), a complex network of proteins and proteoglycans that comprises up to 10% of muscle mass [1]. Muscle regeneration is a tightly regulated process involving intricate cellular and molecular mechanisms. Following injury, matrix metalloproteinases degrade components of the basal lamina, releasing signaling molecules that activate satellite cells. These cells are located between the basal lamina and the apical sarcolemma of myofibers, supported by key ECM components such as laminin and collagen type IV [2, 3]. Activated satellite cells contribute to ECM remodeling by upregulating fibronectin expression and secreting matrix metalloproteinases, particularly MMP-2 and MMP-9. However, the ECM itself is indispensable for facilitating the proper response of satellite cells to these signals [4]. Without intact cell-ECM interactions, even high levels of regulatory factors such as myogenin fail to promote effective muscle differentiation [5]. Moreover, *in vitro* studies demonstrate that satellite cells lose their mitogenic and myogenic potential when removed from their muscle-specific niche [6]. Mechanical properties of the ECM further influence satellite cell behavior and are critical for effective muscle repair, highlighting the importance of biomechanical integrity in the regenerative process [7]. While resident fibroblasts, though limited in number, are the primary contributors to ECM production and assembly, muscle cells also play an active role in ECM remodeling by secreting specific ECM components [8]. Additionally, the immune response is pivotal to muscle repair, particularly in the early stages of injury. Immune cells, such as macrophages, are recruited to the injury site to clear debris, release pro-regenerative cytokines, and support ECM remodeling, thereby facilitating successful regeneration [9, 10].

When muscle damage exceeds the tissue’s regenerative capacity, this innate repair mechanism becomes overwhelmed, leading to volumetric muscle loss (VML) and resulting in permanent functional deficits. Dysregulated or chronic inflammation, often observed in severe VML, exacerbates the hindered muscle regeneration and leads to fibrosis [11, 12]. Despite advances, current clinical approaches to VML treatment face significant limitations. The gold standard, autologous muscle transfer, causes donor site morbidity and is limited to tissue availability. Similarly, cell-based therapies have struggled due to low survival and engraftment rates, limited donor cell availability, *ex vivo*-induced cellular changes, and the risk of teratoma formation with pluripotent stem cells. Acellular scaffolds made of synthetic biomaterials also face challenges due to insufficient cell recognition. Scaffolds composed of individual ECM components like collagen and laminin have shown some potential, but their efficacy remains limited [13–15].

ECM-based acellular products, usually from animal sources but free of antigens and major histocompatibility complexes, have emerged as promising alternatives [16]. Various tissue sources, including skeletal muscle [17, 18], dermis [19], and porcine or bovine bladder [20, 21] and small intestinal submucosa [22, 23], have been explored. However, these scaffolds often lack the ultrastructure and composition specific to muscle tissue, resulting in limited regeneration in VML cases [24, 25]. Recent innovations in more advanced ECM-based therapies for VML aim to address these shortcomings. Hybrid scaffolds that combine synthetic and natural materials are being developed to enhance both mechanical properties and bioactivity [26, 27]. Additionally, techniques such as 3D printing have shown promise in replicating the anisotropic architecture of muscle tissue, which is critical for functional regeneration [28, 29]. The effectiveness of these approaches needs to be seen but overall, acellular therapeutic strategies that promote endogenous cell recruitment represent a promising path forward, offering greater potential for clinical translation. These strategies must replicate the native muscle ECM environment, considering both compositional and structural elements to support tissue regeneration and restore functional recovery in VML [30].

In this study, an Anchored Cell Sheet Engineering platform [31] is utilized to generate completely scaffold- and biomaterial-free, fully formed, and functioning muscle fibers from primary myoblasts. Following decellularization, these fibers served as tissue-specific ECM scaffolds for VML treatment in rat models. Through comprehensive characterization, including proteomics, immunohistochemical analysis, and scanning electron microscopy (SEM), it was demonstrated that these engineered ECM fibers possessed not only appropriate biomechanical properties but also exhibited mature, *in vivo*-like ECM composition and microstructure. To evaluate their regenerative potential, an 8-week study in a rat VML model was conducted, employing multiple assessment methods including histological, immunohistochemical, and functional analyses at various time points. Additionally, a novel image analysis quantification workflow was developed that provided unique insights into the presence and behavior of different cell populations, their contributions to immune and fibrotic responses, and their roles in tissue remodeling and regeneration.

## 2. Results

Tissue-specific skeletal muscle ECM fibers with proper mechanical and microstructural properties, engineered using a scaffold- and biomaterial-free biofabrication platform called Anchored Cell Sheet Engineering, were used to treat VML cases in rat models. The previously published technique [31] (**Figure 1a**) was used for this purpose. The process began by growing multiple layers of primary myoblasts on a patterned PDMS membrane and inducing their differentiation and ECM production over a 14-day period, after which cells and their ECM were easily scraped off the membrane as a coherent sheet. The anchorless sheet recognized the flexible pillars made from Ecoflex 00-30 and utilized them as new anchors to remodel into muscle fibers. This remodeling further enhanced cellular phenotype and ECM content of the fiber, with full maturation achieved by day 18. The muscle fibers were then decellularized and treated with DNase solution to remove cell-free DNA before being used for *in vivo* regeneration of skeletal muscle. The proper ECM composition and microstructure of these fibers were previously demonstrated and are shown here through immunohistochemistry (IHC) and SEM characterization (**Figure 1a**) as well as proteomics analysis (**Table S1 and S2**, **Figure S1 and S2**).

**Figure 1.**
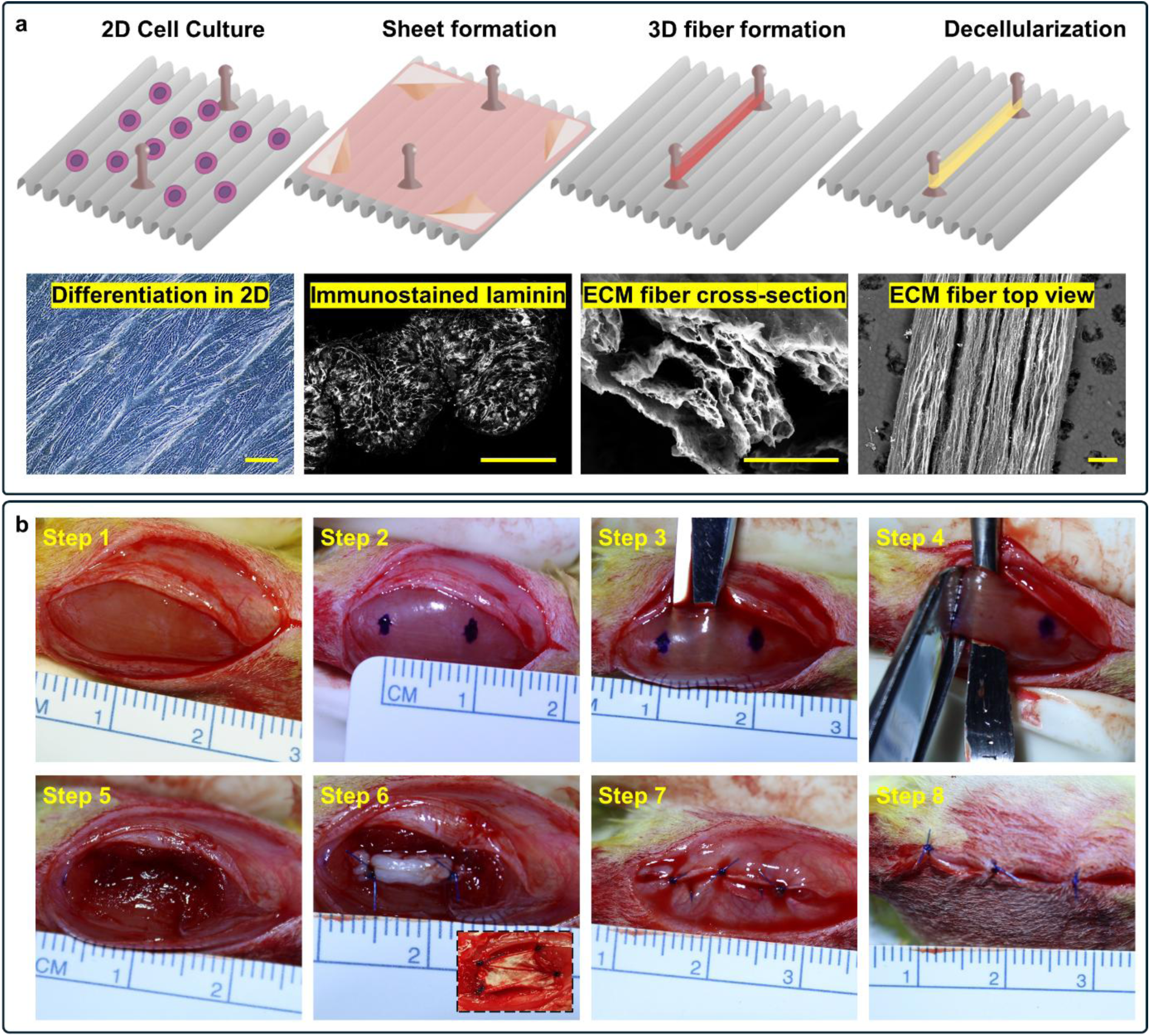
Biofabrication of muscle-specific ECM Fibers and VML surgery protocol. **a)** Schematic representation of the Anchored Cell Sheet Engineering biofabrication process. Primary myoblasts were cultured in multiple layers on patterned PDMS surfaces. Following differentiation and ECM production, the cell sheets were scraped, anchoring to flexible pillars, facilitating further remodeling and maturation of the muscle fibers. These mature fibers were decellularized to leave behind muscle tissue-specific structured ECM. Proper microstructure of the fibers is shown through distribution of laminin as well as aligned components seen in SEM. Scale bars: 200µm. **b)** Surgical procedure for creating VML injury model. The left Tibialis Anterior (TA) muscle was accessed through a lateral fascial incision, and approximately 20% of muscle mass was excised 1 cm from the proximal origin. The defect was treated with either test articles (five acellular ECM fibers, ∼1cm long and ∼1mm thick) or control articles (SIS, shown as insert in Step 6), sutured in place, followed by fascial and skin closure. In the sham group, the defect remained untreated. Engineered ECM fibers can be fabricated with any length to match the defect size, but their fiber-like form factor provides greater flexibility in matching defect shapes by allowing multiple fibers to be implanted in different regions of the VML injury to meet local requirements.

Proteomic analysis revealed the presence of crucial ECM as well as cell adhesion components known to be essential for skeletal muscle function and regeneration. Notable proteins included various collagen types (types I, IV, V, and VI), which provide structural support and mechanical stability. The presence of basement membrane proteins, such as laminins and nidogen, was also confirmed. Additionally, matricellular proteins, including SPARC, thrombospondins (THBS1, THBS2), and tenascin-C, known to modulate cell-ECM interactions and tissue remodeling were detected. The identification of lysyl oxidases and matrix metalloproteinase-2 indicated the preservation of proteins involved in ECM crosslinking and remodeling. Proper microstructure of the fibers was assessed using SEM while proper distribution of ECM components was shown using IHC for laminin. Based on this characterization, the preservation of crucial muscle ECM components and architecture, known to be important for guiding cells in regeneration of muscle *in vivo*, was confirmed and the fibers were used for treating VML.

The 20% VML injuries were surgically created in the Tibialis Anterior (TA) muscle of the left leg in Sprague-Dawley rats by removing 20% of the muscle tissue by weight, 1 cm from its proximal origin. Five acellular ECM fibers (test article) were used to fill the defect site, while commercially available decellularized porcine-derived small intestine submucosa (SIS from Cook® Biotech, control article) was cut to size and used as a control. Both articles were sutured in place. In sham group, the defect was left untreated. The TA muscle epimysium and fascia were then closed. The surgical procedure is illustrated in **Figure 1b**. Animals were euthanized at 2-, 4-, and 8-weeks post-surgery (group sizes detailed in **Table S3**). TA muscles were harvested from both legs, with the contralateral limb serving as the native control. Samples were weighed and fixed for further histological and IHC analysis.

The developments in the damaged muscle tissue were tracked across the three experimental groups, using a novel image processing workflow that assessed the cellular and structural changes, particularly in the actively remodeling site. The workflow began with histology and IHC of tissue samples performed at different time points, followed by whole slide imaging (WSI). WSI files were processed using QuPath^®^ software to manually detect the active site, and for each slide, a high-resolution SVG file was exported containing the contour line defining the region of interest (ROI) and grid lines (500µm apart). A custom Python script was then developed to detect the three layers (original image, ROI contour, and grids) in each SVG file. The script masked each original image by its contour to isolate the ROI and tiled it into smaller pieces of defined sizes using the grid lines (500 µm apart) (**Figure 2a**). For the image analysis purposes, only the middle part of each TA muscle containing the largest cross-section (muscle belly) was utilized. When tile-level data from all biological replicates were used, the assessment covered inter-animal differences and tissue variability within each animal. For tile-level analysis, a mixed-effect statistical model was implemented to account for differences between tiles belonging to different animals. To analyze only the inter-animal differences, slide-level data were used, calculated as the mean value of tiles per slide. For slide-level analysis, a weighted t-test was used to account for the variability inherent to each slide.

**Figure 2.**
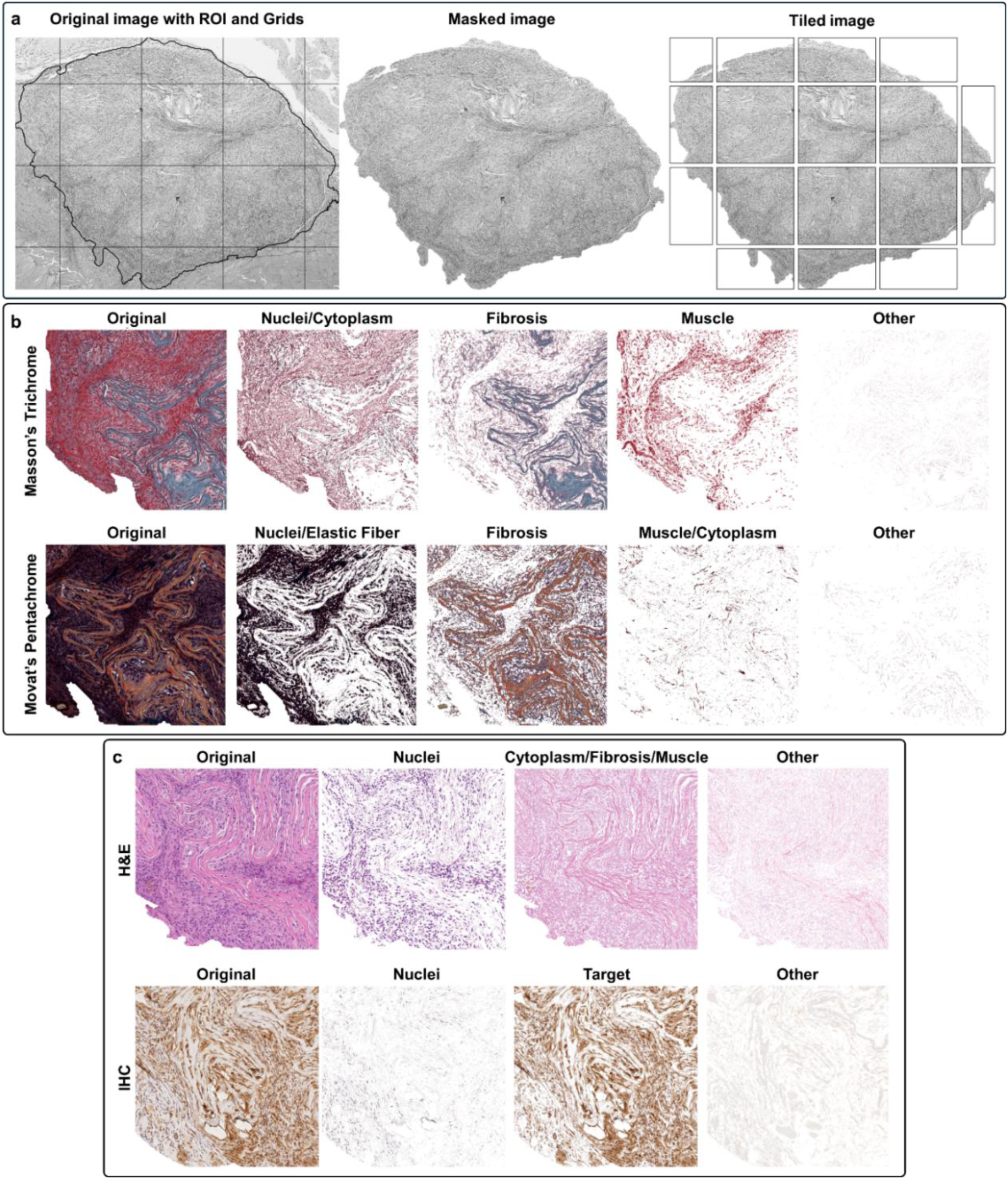
Automated image analysis pipeline for histological assessment. **a)** Workflow schematic showing region of interest (ROI) identification in whole slide images (WSI) using QuPath software. The ROI contour and 500 µm grid overlay were exported as SVG files and processed using a custom Python code to generate masked ROI images and subdivide them into tiles. **b)** Representative segmentation of Masson’s Trichrome (“Nuclei/Cytoplasm” light red, “Fibrosis” blue/green, “Muscle” dark red, “Other” weakly stained) and Movat’s Pentachrome (“Nuclei/Elastic Fiber” dark brown/black, “Fibrosis” yellow/light brown, “Muscle/Cytoplasm” dark red, “Other” weakly stained) staining each with four segments. **c)** Segmentation of H&E (“Nuclei” dark purple, “Cytoplasm/Fibrosis/Muscle”, pink/red, “Other” weakly stained) and IHC (“Nuclei” purple, “Target” brown, “Other” weakly stained) each containing three segments. Area coverage percentage for each segment was calculated per tile. Each complete tile is 500 by 500µm.

The Python script segmented each image based on staining-specific color definitions. Predefined color clusters were established for each staining, and then individual pixels in each tile were assigned to the closest matching cluster. Masson’s Trichrome and Movat’s Pentachrome staining were segmented into four categories (“Nuclei/Cytoplasm”, “Fibrosis”, and “Muscle” for Masson’s and “Nuclei/Elastic Fiber”, “Fibrosis”, and “Muscle/Cytoplasm” for Movat’s), while Hematoxylin and Eosin (H&E) and IHC were divided into three segments (“Nuclei” and “Cytoplasm/Fibrosis/Muscle” for H&E; “Nuclei” and “Target” for IHC) (**Figure 2b and c**). A segment called “Other” was included for all stains that contained weakly stained regions. In H&E stained tiles, nuclei appeared dark purple, while muscle, fibrotic tissue, and cell cytoplasm showed various shades of pink and red, making this stain particularly effective for detecting nuclei. Masson’s Trichrome stained nuclei and cytoplasm light red and muscle dark red, making these components difficult to differentiate reliably. However, its unique blue-to-green staining of fibrotic tissue enabled specific detection of fibrosis independent of composition. Movat’s Pentachrome revealed fibrotic tissue in yellow to light brown, nuclei and elastic fibers in dark brown to black, and muscle and cytoplasm in dark red. In IHC, target antigens appeared light brown and nuclei purple, though intense target staining occasionally masked the nuclei. For quantification, the area coverage percentage was calculated for each segment within individual tiles. After segmentation was performed, random tiles from different staining types were visually assessed to validate the automated segmentation approach. In the current study only the actively remodeling part of the treatment site was considered for quantification as it could be clearly identified with high confidence. Adjacent to this area was the “transition zone” (**Figure S3**) with a combination of native muscle tissue, highly fibrotic tissue, and newly formed myofibers that couldn’t be clearly assigned to intact muscle or treatment area.

The initial assessment of muscle regeneration focused on quantifying the actively remodeling surface area through WSI histology images, measuring total muscle weights across the three time points, and evaluating electrical activity and force generation capacity at the 8-week point in both injured and contralateral muscles. Peripheral tissues including fascia, epimysium, and perimuscular adipose tissues were excluded from the analysis during area analysis, unless these components were integrated into the actively remodeling site. The surface area of the active site was measured using grid lines as reference points by the Python script (**Figure 3a and b**). The active surface area in the test group demonstrated gradual reduction over time, while the control group exhibited accelerated shrinkage with minimal active site by week 8. The sham group showed an even more rapid resolution, with no detectable active site by week 4. These observations were corroborated by muscle weight measurements at week 8, where both sham and control groups displayed significantly lower muscle weights compared to contralateral muscle, while no significant difference was observed between test and native groups (**Figure 3b**). These findings indicated that regeneration had ceased in both sham and control groups by week 8, with no further improvements in muscle regeneration, suggesting that the SIS product had failed to promote VML healing in rats.

**Figure 3.**
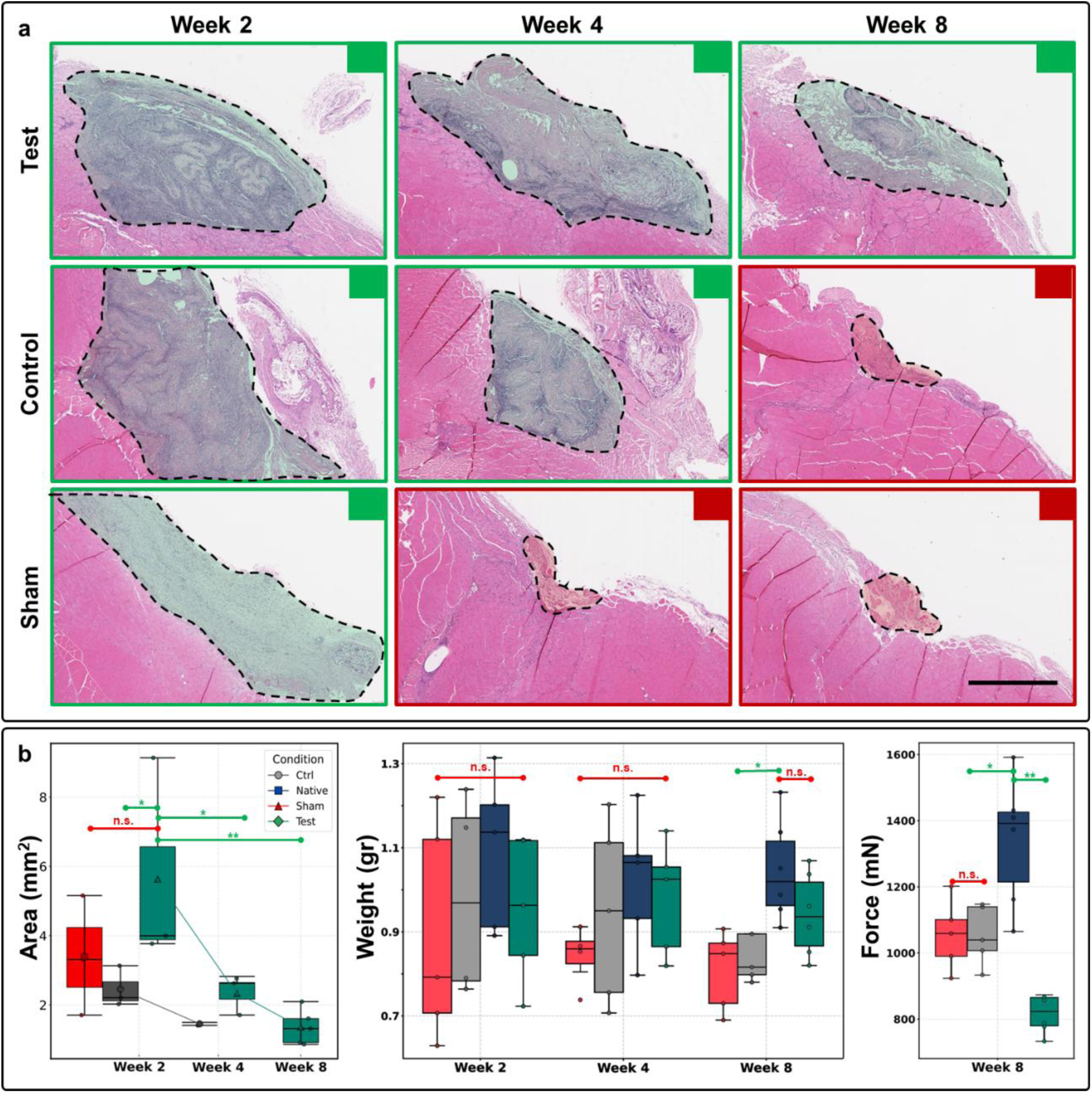
Temporal assessment of VML recovery across treatment groups. **a)** Representative histological sections from the defect center (largest cross-sectional area, muscle belly) showing progression of tissue remodeling. Test group exhibited gradual defect size reduction from weeks 2 to 8. Control group showed rapid initial shrinkage between weeks 2 to 4, with only minimal inflamed tissue remaining at week 8. Sham group displayed no regenerating tissue after week 2, with only residual inflammation visible. Scale bars: 2mm. **b)** Quantitative analysis of tissue recovery. Left: Active site surface area measurements over time (n = 5 per group per timepoint); Center: Muscle weight comparison at different time points, showing 20% reduction in control and sham groups versus native muscle, while test group maintained native-equivalent mass at week 8; Right: *in vivo* functional assessment of muscles by measuring peak isometric force after percutaneously stimulating peroneal nerve. Peak isometric force at maximum stimulation frequency showed 91-92% recovery in sham and control groups versus 77% in test group compared to contralateral muscles. Data are presented as box plots showing median, interquartile range, and individual data points. Lines connect mean values between consecutive time points for each condition. For Area, Weight, and Force measurements across experimental conditions and time points, one-way ANOVA was performed followed by pairwise Welch’s t-tests between conditions within each week. P-value<0.05 was considered significant; n.s. not significant; *<0.05, **<0.01

Functional assessment was performed through *in vivo* measurement of dorsiflexor muscle group contractions induced by percutaneous electrical stimulation of the peroneal nerve. Optimal isometric twitch torque was determined, and peak isometric force was measured in anesthetized rats at the 8-week time point. While sham and control groups demonstrated 91-92% force recovery compared to contralateral legs, the test group showed approximately 77% recovery (**Figure 3b**). Full force measurements at different frequencies are included as **Figure S4**. The lower force generation in the test group was potentially attributed to ongoing active regeneration at the implantation site, suggesting either incomplete force generation capacity or impaired force transmission through the actively remodeling tissue. To determine whether the continued remodeling indicated progressive and slower regeneration or either delayed complete resolution of the test product with no regeneration, or even chronic inflammation, detailed analyses of the cellular response were conducted, as described in subsequent sections.

The nature of cell types populating the acellular ECM fibers in the test group, SIS control, and untreated injury site of the sham, as well as the extent of cellular responses were investigated using histological analysis and IHC for various targets (**Figure 4**), and the quantification workflow, explained in the previous section, was applied to all samples. Histological analysis included H&E, Masson’s Trichrome, and Movat’s Pentachrome (**Figure 5**). In all cases, native muscle tissue from the contralateral leg served as the reference point. These values represented the area coverage of each segment in the WSI images/tiles, excluding the white background. In order to evaluate differences between different parts of each large defect size, tiling was performed to break down each sample to smaller units for analysis. For the statistical analysis, a mixed-effect model was used to account for the presence of multiple tiles from the same animal, as well as for the multiple animals as separate biological replicates.

**Figure 4.**
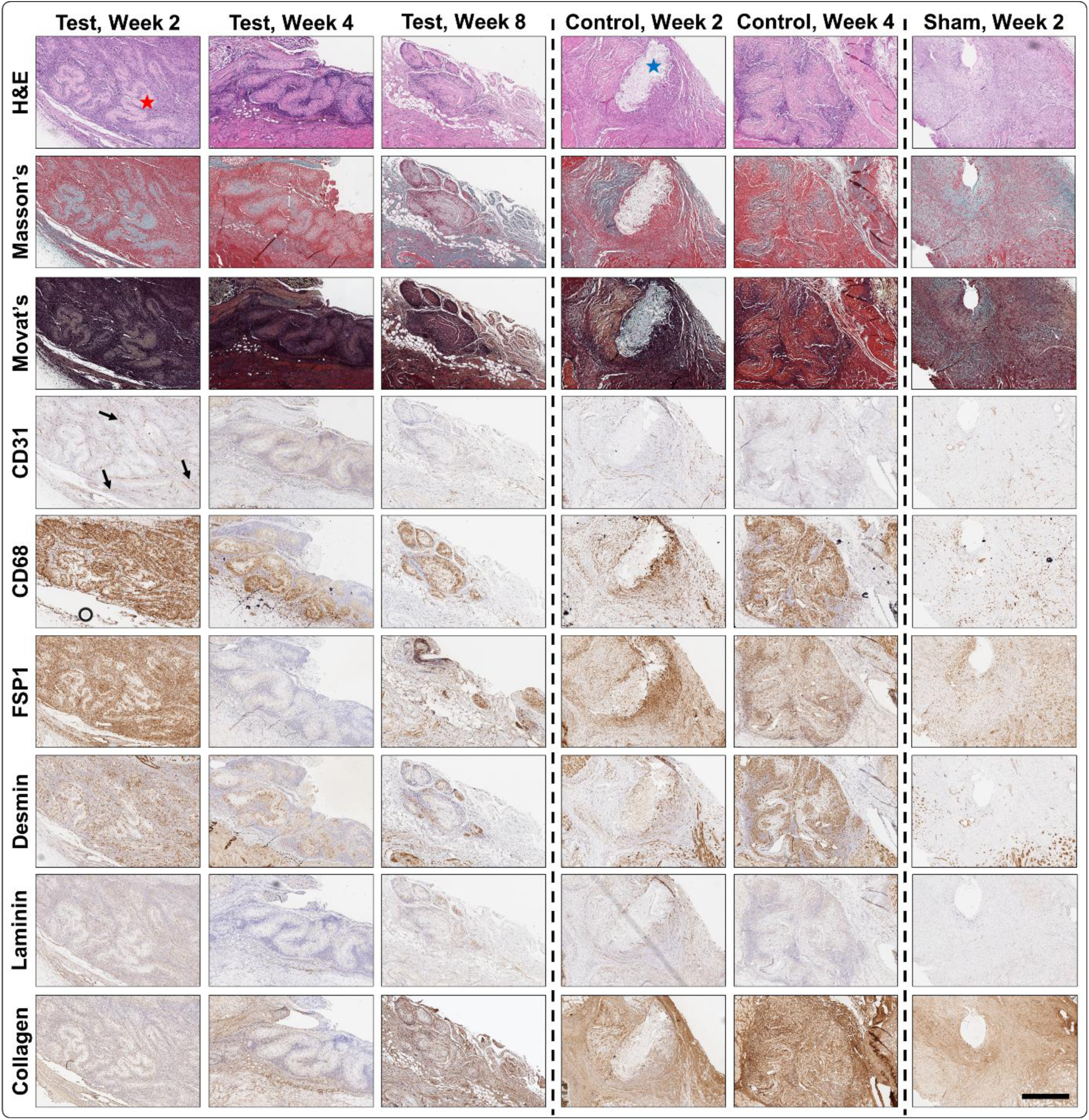
Histology and IHC images of test, control, and sham groups. Temporal changes in the active treatment sites showed distinctive patterns when different staining types and targets were evaluated. These images were further quantified using a combination of QuPath® software and a custom Python script. * shows areas of test article that were not populated by any cell type at the beginning of the process while no such large areas were found in control group. *shows the fat deposition deep into the implanted SIS subject observed early on in some of the control subjects, indicating severe fibrotic response. Not only were high levels of CD31 expression observed in test group, but also lumen-like structures were formed as early as week 2, shown by the arrows. Sham group showed much lower cell densities that were uniformly distributed, mostly fibrotic and inflammatory in nature across the defect area. Scale bar: 2mm.

**Figure 5.**
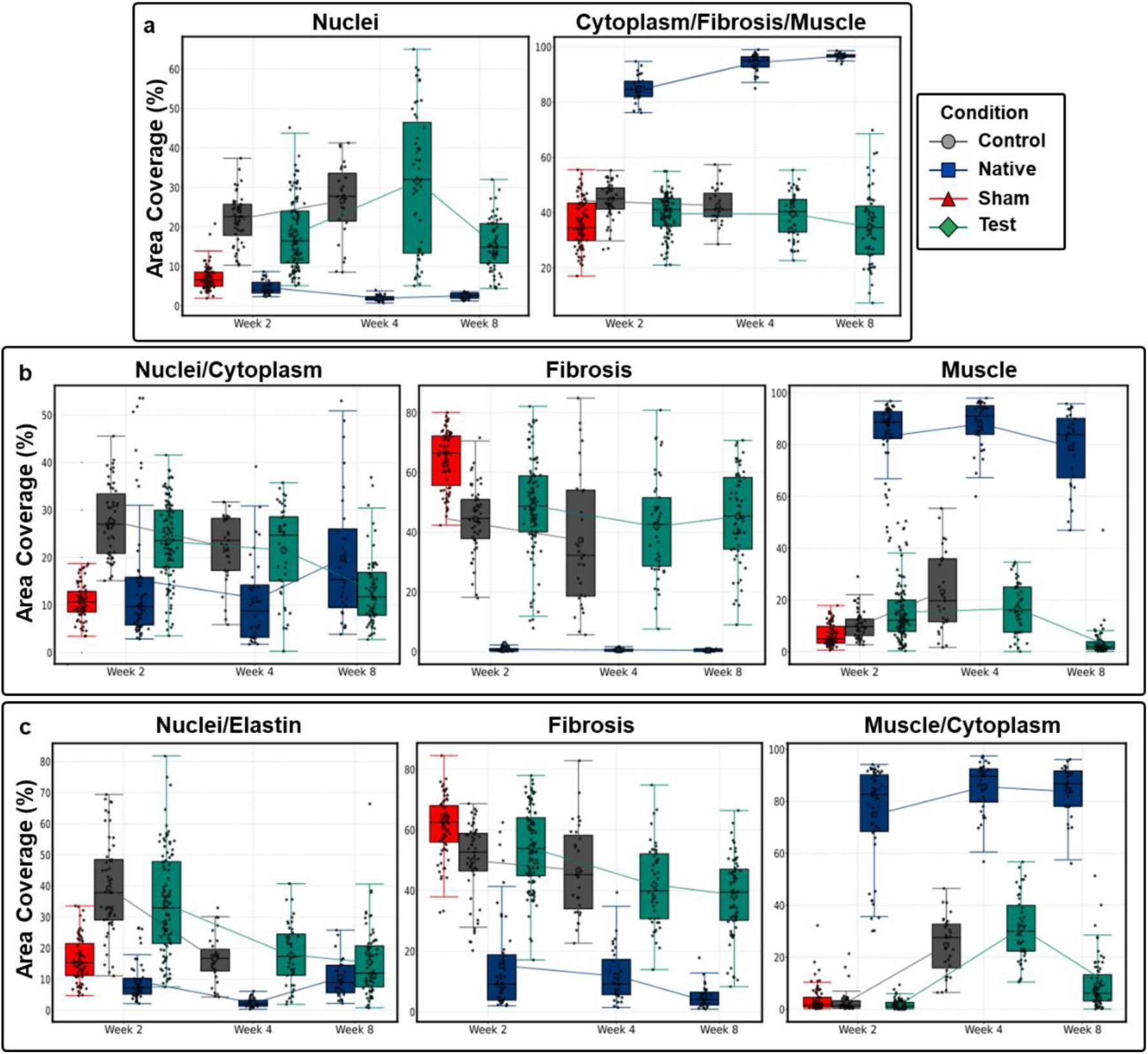
Quantified observations from histology images using tile level data. The percentage area of each segment was recorded across different tiles of histological images from all animals in each condition and time point. **a)** Hematoxylin and Eosin (H&E) staining provided reliable detection of nuclei across all conditions, revealing higher cell numbers in the test and control compared to other groups. **b)** Masson’s Trichrome and **c)** Movat’s Pentachrome staining methods effectively detected fibrotic regions within the active treatment area. The test group exhibited persistent fibrotic tissue presence in both staining groups. Data is presented as box plots showing median (center line), interquartile range (box), and individual data points. Lines connect mean values between consecutive time points for each condition. A mixed-effects model was used for statistical analysis to account for multiple tile measurements within each animal, followed by pairwise Welch’s t-tests between conditions and weeks with Bonferroni correction for multiple comparisons. P-values for all the comparisons are included as heatmaps in **Figure S5**.

Nuclei content analysis using the nuclei segment of H&E staining, which quantified the total number of cells within the active injury site, revealed significantly higher cell counts in both the control and test groups compared to native tissue. In contrast, the cell count in the sham group was much lower, remaining comparable to the native tissue at week 2. This observation indicated that both the test and control articles were successfully populated by host cells; however, nuclei-free regions deeper within the test group (**Figure 4**, indicated by a red star) were identified and further supported by the longer whiskers observed in the H&E nuclei analysis (**Figure 5a**). At week 4, both the test and control groups showed a slight increase in cell numbers, but the test group exhibited a significant decrease by week 8.

Differences between the test and control groups, when compared to both native and sham groups in the Nuclei and Elastic Fiber segment of Movat’s Pentachrome (**Figure 5b**), were more pronounced than those observed in the Nuclei segment of H&E, suggesting a higher elastic fiber content. This finding may indicate greater ECM deposition as early as week 2, though the implanted articles might have contributed to this elastic fiber content. By week 4, these values had decreased, and by week 8, the test group resembled native tissue. The fibrosis segment of Masson’s Trichrome, which included elastic fibers, revealed significantly higher fibrotic tissue deposition in all groups compared to native tissue (**Figure 5a**), with the sham group showing the highest values. In the test group, these fibrosis levels remained consistent across all three time points.

Further analysis of the muscle regeneration process was performed through IHC evaluations of all groups, at both tile- and slide-levels (**Figure 6**). Immune response and presence of fibrotic cells were assessed using CD68 and FSP1 targets, identifying macrophages and fibroblasts, respectively. Vascularization, important for tissue regeneration, was assessed using CD31, while the onset of muscle regeneration was assessed using Desmin as an early marker of muscle differentiation. Laminin content, an important skeletal muscle ECM component involved in guiding skeletal muscle regeneration, and collagen I, an important component of skeletal muscle integrity and a major indicator of fibrosis, were also assessed.

**Figure 6.**
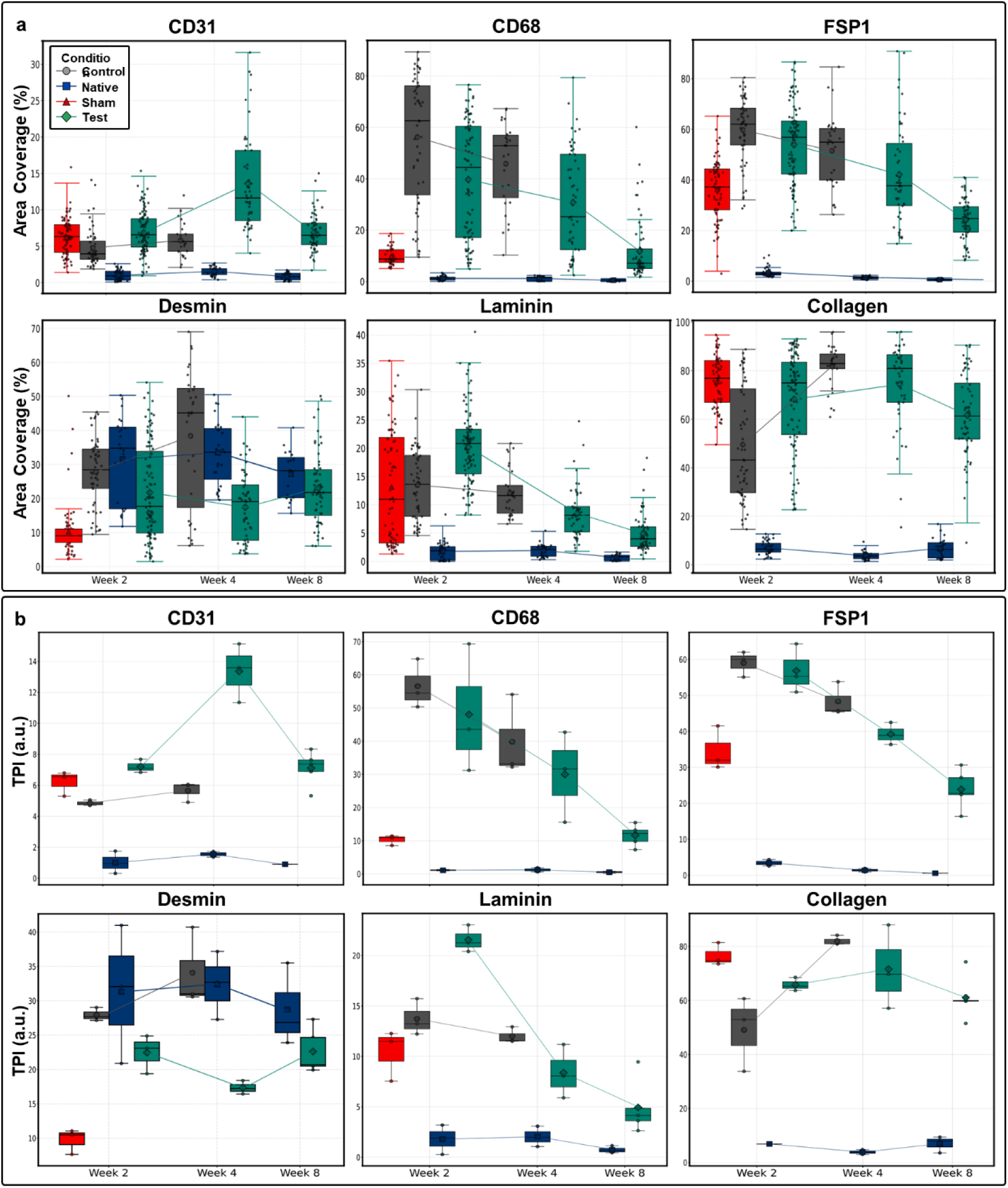
Temporal changes in IHC markers shown **a)** using tile-level data, and **b)** through unitless representation of slide-level data with the Target Prevalence Index (TPI). Various markers for vascularization (CD31), immune response (CD68), fibrotic response (FSP1), muscle regeneration (Desmin), and ECM components (Laminin and collagen I) demonstrated distinct temporal patterns across different treatment groups compared to native tissue. While IHC groups effectively detected specific targets, nuclei detection was compromised in certain cases as target staining occasionally masked the nuclei. TPI was calculated by normalizing the area coverage value for each IHC target to the total nuclei area detected in Hematoxylin and Eosin staining, providing a unitless index for more meaningful temporal comparisons between different conditions and native tissue. The test condition maintained elevated presence of endothelial cells (CD31) and demonstrated progressive formation of vascular lumens over time, while the control group exhibited significantly lower vascularization. Immune cells (CD68) presence gradually decreased to near-native levels in the test group, whereas the control group maintained high levels at weeks 2 and 4. Fibrotic cells (FSP1) demonstrated patterns similar to immune cells in both test and control groups. Desmin index remained lower than native tissue in test group but had inched closer to it by week 8. While collagen levels were sustained in test group, its levels significantly increased in control. Laminin levels showed a marked decrease in test group and only modest decrease in control. Slide level information for each animal was generated and used by calculating mean and standard deviation from all the tiles for that animal. Data is presented as box plots showing median, interquartile range, and individual data points. Lines connect mean values between consecutive time points for each condition. For tile-level and slide-level data a mixed-effect model and a weighted two-sample t-test were used, respectively. P-values for all the comparisons are included as heatmaps in **Figures S6**.

When considering tile-level data (**Figure 6a**), all three groups (test, control, and sham) exhibited elevated levels of CD31-positive cells compared to native muscle. In the control group, these levels remained constant at week 4, whereas the test group showed a spike at week 4, followed by a decrease at week 8, though still remaining higher than native tissue. The sham group displayed a higher presence of immune cells (CD68-positive cells) compared to native muscle; however, its levels were significantly lower than those observed in the test and control groups. In the test and control groups, CD68-positive cell levels remained stable, but the test group showed a decrease by week 8. Fibrotic cell levels (FSP1-positive cells) were significantly elevated in all three groups relative to native tissue, but in the test group, these levels gradually decreased over time, reaching lower levels by week 8. While Desmin levels in the control group remained similar to those of native tissue, suggesting appropriate muscle progenitor recruitment, proper muscle fiber formation was not observed. In contrast, the test group exhibited lower Desmin levels, yet multiple newly formed, isolated, and scattered muscle fibers were detected at week 8 (**Figure 7**). At week 4, the control group displayed a distinct boundary between the implanted SIS and native muscle tissue, whereas the test group showed a transition zone between the native muscle and the ECM fiber implant, characterized by both newly formed myofibers and actively remodeling fibrotic tissue.

**Figure 7.**
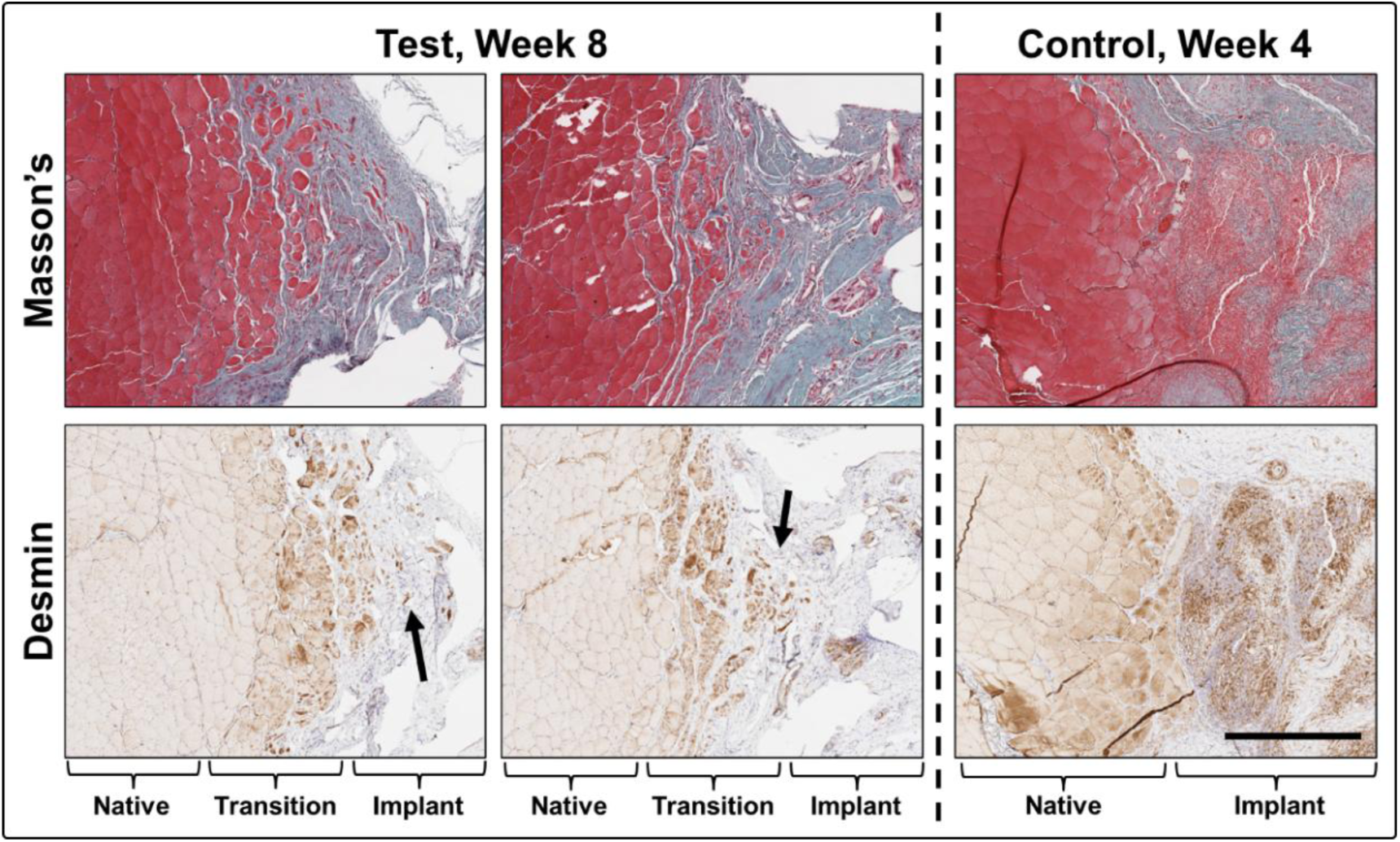
Differences between control and test groups in recruiting muscle progenitor cells and new myofiber formation shown using Desmin IHC staining and Masson’s Trichrome. While quantification at both tile- and slide-level showed more Desmin positive cells in the control group at all time points, a clear transition between native muscle and implanted article was detectable with no proper myofiber formation. At the same time, the test group showed a transition zone where native muscle, the implanted ECM fibers, and both newly formed scattered myofibers (shown by the arrows) and the actively remodeling fibrotic tissue were present. Lack of cohesion between these myofibers could have caused the lower force recorded in the test group. Scale bar: 500µm.

All three groups exhibited significantly higher collagen I content compared to the native tissue. While these levels remained steady between weeks 2 and 4, the test group showed a slight decline by week 8. Given the fibrotic tissue content identified in Masson’s Trichrome staining, this collagen likely originated from both the onset of fibrosis and the composition of the original implanted scaffolds. Additionally, all three groups displayed elevated levels of laminin, a key glycoprotein of the ECM basement membrane. These abnormally high laminin levels may also indicate increased fibrosis. While laminin levels remained high in the control group at week 4, they progressively decreased in the test group from week 2 to week 8, approaching native tissue levels. This decline suggested effective remodeling toward a less fibrotic state in the test group.

Overall, the results indicated that the initial response in all three groups was predominantly fibrotic and inflammatory. In the sham group, fewer cells were present at week 2, as expected due to the absence of any 3D scaffolding support. The control group exhibited sustained levels of high immune activity, fibrosis, and ECM content across the first two time points; however, the lack of both new myofiber formation and recovery of tissue weight suggested that this heightened cellular activity was detrimental and counterproductive to the muscle regeneration process. In contrast, only the test group was able to maintain muscle tissue volume and weight while displaying more moderate cellular responses, with both immune and fibrotic activities decreasing over time. This group also demonstrated a gradual remodeling of the ECM fibers and a progressive reduction in fibrotic ECM content. Proper vascularization, essential for natural healing, was observed in the test group, along with recruitment of Desmin-positive cells at levels comparable to native tissue. However, the formation of only a few scattered myofibers suggested a potential delay in the regeneration and maturation of fully developed muscle tissue, possibly requiring immune and fibrotic responses to subside further. These findings underscore the importance of longer-term studies to capture the full regenerative potential of the engineered ECM fibers, supported by the preservation of muscle volume and weight in the test group.

A unitless index, termed the Target Prevalence Index (TPI), was defined for IHC targets to complement the area percentage comparisons presented earlier (**Figure 6b**). This index was calculated using slide-level data (mean values across all tiles for each animal) rather than tile-level data to facilitate a clearer comparison of overall changes across different conditions over time. To normalize target expression relative to total cell count, linear regression was employed, with the percentage of nuclei area (from H&E staining) as the independent variable and the percentage of target-positive area (from IHC staining) as the dependent variable. This normalization method was chosen over simpler approaches, such as directly dividing values, to avoid compounding uncertainties (standard deviations) from the two measurements, which could artificially inflate the variability of the final index values. The final TPI values, derived from regression residuals centered around the original target expression means, reflected the deviation of actual target expression from expected values based on cell density. This allowed for an accurate quantification of target prevalence while accounting for variations in cell density. The approach effectively normalized IHC data for both cellular (CD31, CD68, FSP1, and Desmin) and extracellular (Laminin and Collagen) targets.

The slide-level TPI enabled normalization of each target’s expression relative to total cell count, facilitating meaningful comparisons across conditions. While the TPI values exhibited trends similar to those observed in direct area comparisons, the distinctions between conditions over time became more pronounced. In the test group, TPI analysis revealed a sharp decline in CD68- and FSP1-positive cells, as well as Laminin levels, though all three remained elevated compared to native muscle, reflecting an active but rapidly subsiding fibrotic and inflammatory responses. Desmin TPI in the test group was initially lower than in native tissue, experiencing a slight decrease at week 4 before increasing by week 8, approaching native tissue levels. Notably, the control group displayed higher Desmin TPI values than both the test group and native tissue but lacked any evidence of new myofiber formation (**Figure 7**). During the first two weeks, the control group also exhibited elevated CD68 and FSP1 levels compared to the test group, accompanied by a marked increase in collagen deposition. These findings, combined with lower CD31 TPI values, suggested a heightened fibrotic and inflammatory response with low vascularization, creating a less favorable environment for muscle regeneration and healing. In the sham group, TPI indices indicated a predominantly fibrotic response rather than a significant inflammatory one, further highlighting differences in the healing dynamics among the three groups.

## 3. Discussion

Treatment of VML, among other severe damages to skeletal muscle tissue, poses a significant challenge in regenerative medicine due to the complexities of skeletal muscle structure and function. With no existing standard of care for VML, no tissue engineering or regenerative medicine approach has shown proper restoration of function in injured skeletal muscles yet [29]. Here, engineered skeletal muscle -specific ECM fibers, fabricated using Anchored Cell Sheet Engineering, a scaffold-free biofabrication platform capable of recreating physiologically relevant ECM composition *in vitro* entirely by the cells, holds considerable promise as a therapeutic strategy for VML. This innovative method addresses several limitations of conventional ECM scaffolds, offering a new approach for muscle regeneration and providing a foundation for advancing VML treatment.

One of the most common treatments used for treating VML is animal-derived decellularized ECM from non-muscle tissues such as SIS and UBM. While these materials, among other acellular scaffolds, have shown efficacy in some tissue engineering applications, their capacity to support skeletal muscle regeneration has been limited [32, 33]. These animal-derived acellular ECM products often lack the mechanical robustness needed in such load-bearing tissues, and their composition does not match the intricate skeletal muscle tissue ECM [24, 25]. It’s been shown that use of animal-derived ECM in VML injury models yields insufficient muscle fiber regeneration as these scaffolds are often remodeled rapidly at the injury site and only lead to high levels of fibrotic tissue formation, inhibiting any chances of muscle regeneration [32, 34]. In cases of more complex musculoskeletal trauma, the use of animal-derived ECMs such as SIS has been shown to impair healing of the adjacent bone as well [23].

It is known that tissue specific microenvironments, including proper biochemical and biophysical properties, are necessary for guiding cellular behavior to promote tissue organization. This is key for effective tissue engineering and regenerative medicine of skeletal muscle tissue [35, 36]. Therefore, it is reasonable to speculate that matching the acellular ECM composition and microstructure to that of skeletal muscle tissue is of paramount importance for guiding host cells toward its regeneration. Proteomic analysis of the engineered ECM fibers in this study revealed that they closely mimicked native skeletal muscle ECM, including key structural proteins (collagen types I, IV, V, and VI), basement membrane proteins (laminins and nidogen), and matricellular proteins (SPARC and thrombospondins) (**Table S1 and S2**, **Figure S1 and S2**). IHC for laminin showed proper distribution of this key ECM protein in guiding regeneration of muscle (**Figure 1a)**. This is particularly important as, during myogenesis, Laminin enhances myoblasts proliferation and migration, and later their alignment before fusion [37].

SEM images also showed the proper alignment and directionality of muscle ECM as well as proper microstructure including its porosity (**Figure 1a**), all known to be of utmost importance in guiding cellular behavior during muscle healing and regeneration. The fiber-like form factor of the engineered ECM fibers in this study, with their tunable lengths, provides greater flexibility to match local defect shapes. Multiple fibers of different lengths can be implanted at various locations within the muscle to better align with muscle orientation requirements. This advantage is particularly important since mechanical manipulations (such as folding and securing) of sheet-like products, including animal-derived ECMs, can disrupt their integrity and compromise their biological activity when fitted to the complex geometry of VML injuries, potentially leading to more variable functional outcomes [38].

The use of decellularized animal-derived ECM scaffolds for treating VML has been hindered by several other factors related to the decellularization process itself. While decellularization aims to remove cellular components, residual cell-free DNA often remains in the scaffolds, potentially triggering inflammatory responses and complicating the regenerative process. Moreover, the strong detergents required to effectively decellularize thick and dense animal tissues can leave behind residues that are toxic and induce foreign body responses. These detergent residues have been shown to increase inflammatory markers like IL-1β and impede cell infiltration in animal models. The harsh decellularization methods necessary for complete cell removal can also damage the ECM itself, altering its composition, structure, and bioactivity [39–41]. These have the potential to compromise the ECM’s ability to provide appropriate biochemical and biophysical cues that can impede cell’s natural processes essential for muscle regeneration in VML cases. It was shown that co-delivery of decellularized UBM with autologous minced muscle even negatively affected the regenerative capacity of the autografts in a rodent model of VML [42]. Meanwhile, the *in vitro* engineered ECM fibers in the current study, while thinner and not as dense as animal derived tissues, were decellularized individually using less harsh detergents, showing below toxic levels of both detergents and cell-free DNA (data not shown) contributing to the less severe inflammatory and fibrotic responses observed in the rat models (**Figure 5 and 6**).

The effectiveness of the engineered ECM fibers in treating VML was assessed through histological and IHC evaluations, comparing them against a commercial SIS product as the control and a sham group with no intervention. The quantitative analysis of these images presented unique challenges that required development of a novel workflow. Traditional approaches often lack standardization and can be subject to observer bias. The methodology presented here, combining QuPath® software for ROI detection with custom Python scripts for automated segmentation based on staining-specific color definitions, offered a more objective and reproducible approach to tissue analysis. The systematic tiling of samples and careful consideration of segment categories for different stain types enabled consistent quantification and proper insight extraction across all samples. Importantly, the analysis incorporated proper statistical analysis in each case, for example a mixed-effects statistical model to account for both inter-animal variability (differences between animals) and intra-animal variability (differences between different segments of the treatment area in each animal) when tile level data was used. This hierarchical approach to data analysis provided a more accurate representation of biological variation and strengthened the statistical validity of the findings compared to traditional methods that might overlook such nested structures in the data. The combination of standardized image processing and robust statistical analysis provided a framework that can be valuable for future studies in the field, particularly when dealing with complex tissue regeneration processes that exhibit high biological variability. This workflow, optimized for analyzing the spatial heterogeneity of muscle regeneration in VML models, can also be implemented to other *in vitro* and *in vivo* applications.

The temporal analysis of cellular responses underscored the benefits of tissue-specific ECM in modulating the inflammatory and fibrotic processes critical to tissue remodeling. For image analysis, only the largest cross-sections, representing areas with the greatest mass transfer limitations, were evaluated. Both animal-derived ECM and *in vitro*-engineered ECM fibers demonstrated proper integration with host tissues; however, some subjects in the control group exhibited fat deposition at the interface between the muscle tissue and the implanted material (**Figure 4**). Such fibro-fatty tissue replacements are commonly observed in cases of repeated acute injuries, extensive volumetric muscle loss, or chronic muscle damage, where complete muscle repair is not achievable [9]. In the test group, integration was more robust, as evidenced by the presence of a transition zone containing both newly formed myofibers and an actively remodeling implant site. In contrast, the control group exhibited a distinct boundary separating the native muscle from the implanted material (**Figure 7**).

Both the engineered ECM fibers and the SIS control elicited an initial immune response characterized by CD68-positive cells. However, the engineered ECM fibers demonstrated a controlled reduction in inflammation by week 8 (**Figure 6**). Acute immune responses triggering excessive inflammation have been shown to hinder repair and regeneration [43–45]. The high immune activity observed in the control group may explain the complete degradation of the SIS by week 4, whereas the engineered ECM fibers persisted beyond 8 weeks, supported by a more moderate immune response. The vascularization dynamics further underscored the therapeutic potential of the engineered ECM fibers. Elevated levels of CD31-positive cells (**Figures 6**), along with evidence of lumen formation (**Figure 4**), suggested a well-regulated angiogenic process, in stark contrast to the erratic vascularization patterns often seen with non-specific ECM materials [36, 46]. The fibrotic response, assessed through FSP1-positive cell counts, was also better controlled in the engineered ECM fibers, with levels decreasing over time. While early fibrotic tissue deposition is essential for maintaining tissue integrity in large injuries, persistent fibrosis, often a drawback of non-specific ECM scaffolds, can impair regeneration. This was mitigated in the engineered ECM fibers, likely due to the muscle-specific ECM’s tailored biomechanical and biochemical properties. These findings highlighted a balanced tissue remodeling process conducive to functional regeneration [26, 47]. The introduction of the TPI index, a normalized and unitless metric, further streamlined the comparison of various IHC targets, enhancing the robustness of the analysis.

Force recovery in the engineered ECM fibers was lower (77%) than in the control group (91%), but this discrepancy may reflect active muscle regeneration rather than premature fibrosis. This hypothesis was supported by the formation of scattered new myofibers within a transition zone, indicative of ongoing integration with native tissue (**Figure 7**). Rapid force recovery observed in animal-derived acellular ECM scaffolds has been associated with non-functional fibrotic tissue, providing a connective tissue bridge between damaged segments of skeletal muscle rather than true muscle regeneration. At the optimal muscle length, this bridge can accommodate improved muscle fiber shortening and enhanced contraction in the presence of minimal muscle regeneration [48–51]. For example, in a rodent VML model using porcine UBM, significantly fewer myosin-positive fibers were formed compared to autografts, despite higher initial force recovery. Human clinical studies have also revealed the complexity of force recovery in VML. In a trial involving 13 patients with varying muscle injuries, implantation of porcine SIS ECM resulted in an average strength improvement of 37.3% at six months. However, the study noted that these strength gains did not directly correlate with muscle regeneration [36, 52]. It’s been also shown that animal-derived ECM scaffolds can facilitate new myofiber formation only when combined with muscle stem cells [53]. In contrast, the engineered ECM fibers in this study preserved muscle volume and weight and showed new myofiber formation suggesting a regenerative process that balanced early remodeling with long-term functional outcomes. The regenerative capacity of the engineered ECM fibers can further be improved by addition of drug/gene-releasing mechanisms [54, 55].

To fully validate these findings, further studies are needed to optimize the clinical applicability of this approach and assess its long-term effects. While this study’s eight-week observation period provided valuable insights, it also highlighted the need for extended evaluations to capture the full regenerative timeline. The delayed presence of Desmin-positive cells, forming only scattered myofibers, points to the possibility of a more gradual regenerative process. Despite these limitations, the findings establish a strong foundation for the use of engineered tissue-specific ECM fibers in VML treatment. The observed balance of immune modulation, vascularization, muscle volume preservation, and weight maintenance reflected a physiologically relevant regenerative process, distinguishing this approach from traditional non-specific ECM scaffolds. With further refinement and long-term studies, this scaffold-free biofabrication technique has the potential to revolutionize VML treatment and improve outcomes for patients facing this challenging condition.

## 4. Methods

### Biofabrication of skeletal muscle-specific ECM fibers

Acellular skeletal muscle-specific ECM fibers were fabricated using a previously published technique called Anchored Cell Sheet Engineering [31]. Briefly, custom culture devices were fabricated using multiple resin types, with polydimethylsiloxane (PDMS) (Dow, SYLGARD™ 184) serving as the base membrane for two-dimensional (2D) cell culture, Ecoflex™ 00-30 (SMOOTH-ON) used for the pillars, and Ecoflex™ 00-50 (SMOOTH-ON) used for device walls. Master molds for these devices were made using fused deposition modeling (FDM) 3D printing with acrylonitrile butadiene styrene (ABS) filament. The surface patterns from the 3D-printed molds were replicated onto the PDMS membrane, facilitating cell alignment and enhancing ECM production. The PDMS membranes were treated with tannic acid (Sigma Aldrich, 403040) to improve cell attachment. Primary rabbit skeletal muscle cells (Sigma Aldrich, RB150-05) were cultured on the PDMS culture devices over 18 days, during which cells were added four times (on days 1, 4, 7, and 10) to create a multi-layer cell culture system. The growth medium (Sigma Aldrich, RB151-500) was replaced with differentiation medium (Sigma Aldrich, 151D-250) on day 3, which was refreshed every other day. The differentiation medium was supplemented with 100 µg/mL 2-Phospho-L-ascorbic acid trisodium (Sigma Aldrich, 49752) to stimulate ECM production. On day 14, the multi-layered cell constructs were gently scraped from the edges of the culture device using a sterile cell scraper, facilitating the formation of cell sheets. These sheets were then carefully detached from the membrane and anchored themselves between the two pillars, transitioning to a three-dimensional (3D) culture system. Over the next 4 days, the anchored cell sheets underwent remodeling, resulting in the formation of mature, aligned muscle fibers, rich in mature muscle-specific ECM.

The mature fibers were subsequently decellularized using a detergent solution containing 0.2% v/v Triton X-100 (Sigma Aldrich, 93443) in Hank’s Balanced Salt Solution (HBSS) (Sigma Aldrich, H9269). The decellularization process was carried out over 4 days at 4°C, with the detergent solution being refreshed once at the midpoint of the process. Following decellularization, the acellular ECM fibers were thoroughly washed multiple times with phosphate-buffered saline (PBS) (Sigma Aldrich, P4474) to remove any detergent residues. The ECM fibers were then treated with a 1:1000 dilution DNase solution (Thermo Scientific™, FEREN0521) at room temperature for 1 hour to eliminate any remaining DNA. After DNase treatment, several more additional washing steps with PBS was performed. All procedures were performed inside a biosafety cabinet to maintain sterility. Finally, the decellularized ECM fibers were submerged in PBS containing 1% penicillin-streptomycin, frozen at -80°C, and maintained under these conditions until required for surgery.

### Animal study ethics approval

Animal experiments were conducted by Labcorp (Bedford, USA), an independent contract research organization. Following initial safety assessments, the full study protocol was reviewed and approved by the Institutional Animal Care and Use Committee (IACUC) (protocol number: 2024-NR-09). All animal procedures were performed in accordance with the NIH Guide for the Care and Use of Laboratory Animals. Animals were housed in a temperature-controlled environment (22 ± 2°C) with a 12-hour light/dark cycle and provided ad libitum access to standard rat chow and water. All efforts were made to minimize animal suffering and reduce the number of animals used.

### Surgical model and experimental design

Sprague-Dawley rats received appropriate doses of analgesia before anesthesia. Each rat’s left pelvic limb underwent aseptic preparation and draping. A lateral incision was created from the knee to ankle level, and the tibialis anterior (TA) muscle was exposed through blunt dissection. The TA fascia was incised and carefully separated from the muscle belly. Using a scalpel blade, the central 10-15 mm portion of the TA muscle was marked at its proximal and distal boundaries. Approximately 20% of the muscle mass was removed by sharp dissection, performed by holding the proximal end of the incised tissue and reflecting it distally while cutting the base with a scalpel. The excised muscle tissue was weighed to confirm the percentage of TA muscle removed. The defect was filled with five engineered ECM fibers (test article), which were secured to the remaining TA muscle using polypropylene suture at the corners and margins of the implant. For the control group, porcine SIS ECM (Cook Myotech™, G12581) was trimmed to match the defect size and was sutured following the same technique. The sham group received no treatment in the defect area. The sutures, which incorporated the muscle epimysium to ensure better retention, also served as markers for the defect-implant interface during harvest. The TA muscle fascia was sutured using 5-0 Vicryl in a simple interrupted pattern, followed by skin closure using 5-0 Prolene in the same pattern. During recovery, a compression bandage was applied to the leg for 5-10 minutes. At 2-, 4-, and 8-weeks post-surgery, animals were weighed and euthanized via CO2 inhalation. TA muscles were harvested from both legs, with the contralateral leg serving as an internal control. After weighing, the dissected muscles were placed in 10% neutral buffered formalin and kept at room temperature for further analyses. (details of animals numbers per group per time point in **Table S3**)

### Histological and Immunohistochemical Analyses

The fixed tissue samples underwent dehydration through a series of graded ethanol solutions, followed by xylene clearing and paraffin embedding. Using a rotary microtome, 5 μm thick sections were prepared and mounted onto positively charged glass slides. The samples were processed following standard protocols for Hematoxylin and Eosin (H&E), Masson’s trichrome, and Movat’s Pentachrome staining. For immunohistochemistry (IHC) procedures against CD31, CD68, FSP1, Desmin, Laminin, and Collagen type I, the sections were first deparaffinized using xylene and rehydrated through decreasing concentrations of ethanol. Antigen retrieval was conducted using citrate buffer (pH 6.0) in a pressure cooker for 20 minutes, followed by a 10-minute treatment with 0.3% hydrogen peroxide to neutralize endogenous peroxidase activity. The sections underwent blocking with 10% donkey serum in PBS for 1 hour at room temperature. Primary antibody application was performed overnight in a humidified chamber at 4°C. Following PBS washing, the sections were treated with peroxidase conjugated donkey anti mouse secondary antibody (Jackson ImmunoResearch, 715-035-150) or peroxidase conjugated donkey anti rabbit secondary antibody (Jackson ImmunoResearch, 711-035-152). The complete antibody list is provided in **Table S4**. DAB chromogen (Vector Laboratories) was used to visualize the immunoreaction. After hematoxylin counterstaining, the sections were dehydrated, cleared, and sealed with permanent mounting medium. All stained slides were digitally scanned at 40× magnification using a Leica Aperio AT2 scanner (Leica Biosystems, Buffalo Grove, IL). Image analysis and quantification were performed using Aperio ImageScope software (version 12.4, Leica Biosystems).

### Image Analysis and Quantification

Whole slide images (WSIs) of stained tissue sections were analyzed using a systematic, multi-step computational approach. Initially, regions of interest (ROIs) comprising only the actively remodeling injury/treatment sites were identified in QuPath® software using semi-automated boundary detection based on tissue morphology and architecture, and were marked using a contour line. Reference grid lines were overlaid at 500µm intervals for standardization of spatial measurements across samples. These three layers were then saved in SVG format to preserve imaging details. A custom image processing pipeline was developed and implemented in Python to analyze these WSIs. The code first detected the three layers from each SVG file and saved them individually as high-resolution (300 dpi) PNG files. It then isolated each ROI from the entire slide using binary masks created from the contour line, excluding surrounding regions. The masked regions were then subdivided into analysis tiles using the reference grid lines detected through adaptive thresholding and Hough transform techniques. Each tile maintained experimental traceability through a comprehensive naming convention incorporating treatment condition (Test, Sham, or Control), time point (Week 2, 4, or 8), staining type, and animal number. Empty tiles were excluded from analysis. In the next phase of the process, tissue components defined by the staining type were quantified using a color-based segmentation approach specific to each staining type. For histological stains, H&E (“Nuclei” dark purple, “Cytoplasm/Fibrosis/Muscle” pink/red, “Other” weakly stained), Masson’s Trichrome (“Nuclei/Cytoplasm” light red, “Fibrosis” blue/green, “Muscle” dark red, “Other” weakly stained), and Movat’s Pentachrome (“Nuclei/Elastic Fiber” dark brown/black, “Fibrosis” yellow/light brown, “Muscle/Cytoplasm” dark red, “Other” weakly stained) staining, tissue components were categorized based on their characteristic colors. For IHC targets (“Nuclei” purple, “Target” brown, “Other” weakly stained), the analysis distinguished between DAB-positive regions (brown) and negative regions (purple counterstain). Segmentation was performed using a nearest-neighbor classification in RGB color space, with predefined color clusters established through comprehensive analysis of representative images. The area coverage for the stained part of each tile was measured, and the values were recorded for each condition at each time point. The distribution of tissue components across conditions and time points was visualized using box plots with median (horizontal line), interquartile range (box), and min/max values (whiskers) for each anatomical location. For statistical analysis, a mixed-effects model was employed to account for both biological replicates (different animals) and technical replicates (multiple tiles per slide from each animal). This was then followed by a pairwise t-tests with unequal variance assumption (Welch’s t-test) between groups at different timepoint. P-values were adjusted for multiple comparisons using the Bonferroni correction method and values less than 0.05 were considered statistically significant.

To provide a more meaningful temporal comparison between different conditions an index called Target Prevalence Index (TPI) was defined for IHC targets by fitting a linear regression between the percentage of nuclei area from H&E staining as the independent variable and the percentage of IHC target-positive area as the dependent variable. The TPI values, derived from regression residuals normalized to the original target expression means, provided a unitless measure of target abundance independent of local cell density. TPI values were presented using the same box plots. For calculating TPI values, slide level data (mean of all tile values for each slide) were used with a weighted two-sample t-tests with Bonferroni correction implemented for statistical analysis purposes to account for standard deviations calculated from tile level data.

### Neuromuscular Function Assessment

Muscle performance was measured in vivo at week 8 with a 305C muscle lever system (Aurora Scientific Inc., Aurora, CAN). Rats were anesthetized with isoflurane (2% induction, 2% maintenance) and placed on a thermostatically controlled platform at 37°C. The knee was pinned using a 25G needle pushed against the proximal tibia and secured to a stabilization device, with the foot firmly fixed to a footplate on the motor shaft. Contractions of dorsiflexor muscle group were elicited by percutaneous electrical stimulation of the peroneal nerve and optimal isometric twitch torque determined by measuring responses at increasing stimulation intensities. A series of stimulations were then performed at increasing frequency of stimulation (1-150 Hz, 0.2ms pulse width, 500ms train duration) and maximal peak isometric force determined at each frequency. Maximum force values were visualized using box plots, and force-frequency relationships were plotted for each experimental group. The method follows the protocol provided in reference [56]. Statistical analysis was performed using one-way ANOVA followed by pairwise t-tests with unequal variance assumption (Welch’s t-test) to compare differences between groups. P-values less than 0.05 were considered statistically significant.

## 5. Conclusions

This study established engineered tissue-specific ECM fibers, fabricated *in vitro* using a completely scaffold- and biomaterial-free platform, as a promising therapeutic strategy for VML, offering significant advantages over traditional non-specific and often animal derived ECM scaffolds. The fibers demonstrated the ability to preserve muscle volume and weight while promoting controlled immune and fibrotic responses and vascularization. Unlike conventional animal-derived ECM scaffolds, which are prone to rapid degradation and excessive and chronic inflammation and fibrosis, the engineered fibers exhibited a tailored regenerative process with progressively reduced fibrosis and sustained tissue remodeling. The use of a gentler and more effective decellularization process preserved the biochemical and structural integrity of the ECM, minimizing inflammatory complications commonly associated with traditional animal-derived ECMs. Although the use of engineered ECM fibers led to the formation of scattered myofibers, force recovery was slightly lower than the control group. Histological and immunohistochemical findings suggested that this discrepancy reflected active regeneration rather than impaired healing, highlighting the potential for sustained long-term functional recovery. Furthermore, the novel image analysis workflow developed in this study offered a reproducible and objective framework for assessing spatial tissue heterogeneity, advancing the field of regenerative medicine in general. While the eight-week timeline provided valuable insights, extending the observation period is critical to fully elucidate the regenerative potential of these engineered fibers. With continued refinement and longer-term studies, this scaffold-free biofabrication approach holds the potential to transform VML treatment, paving the way for improved outcomes in patients with severe muscle injuries.

## Acknowledgment

The authors expressed gratitude to Caroline O’Neil from Robarts Research Institute, Western University, London, Canada for performing the histology assessment; and Jennifer Martin and Ramzi Khairallah from Myologica, Baltimore, United States for muscle performance assessments. All other experiments were performed at Velocity Incubator, Waterloo, Canada. The authors appreciate the support from Velocity Incubator’s staff members.

## Data Availability Statement

The custom Python script used in analyzing the data is available on <u>GitHub</u>. Any other data used in the study but not included here is available upon request.

## Declaration of use of generative AI

During the preparation of this work, the authors used Open AI’s ChatGPT and Anthropic’s Claude in order to grammatically edit the manuscript and debug the Python scripts. After using these tools, the authors reviewed and edited the content as needed and take full responsibility for the content accuracy.

