## Supplementary Information for "Muscle-Specific ECM Fibers Made with Anchored Cell Sheet Engineering Support Tissue Regeneration in Rat Models of Volumetric Muscle Loss"

**Table S1.** List of extracellular matrix proteins identified in acellular fibers using proteomics analysis, generated using the STRING database.

| **Detected proteins** | |
|---|---|
| SERPINF1 | Serpin family F member 1; Belongs to the serpin family. (418 aa) |
| COL6A2 | Uncharacterized protein. (425 aa) |
| LOXL1 | Lysyl oxidase like 1. (535 aa) |
| VCAN | Versican. (3473 aa) |
| CCN2 | Cellular communication network factor 2. (437 aa) |
| GLG1 | Golgi glycoprotein 1. (1225 aa) |
| THBS1 | Thrombospondin 1. (1171 aa) |
| FBN1 | Fibrillin 1. (2831 aa) |
| LUM | Lumican. (338 aa) |
| EMILIN1 | Elastin microfibril interfacer 1. (982 aa) |
| COL6A3 | Uncharacterized protein. (562 aa) |
| FBN2 | Fibrillin 2. (2912 aa) |
| ECM2 | Extracellular matrix protein 2. (691 aa) |
| TIMP3 | Metalloproteinase inhibitor 3; Complexes with metalloproteinases (such as collagenases) and irreversibly inactivates them by binding to their catalytic zinc cofactor. May form part of a tissue-specific acute response to remodeling stimuli (By similarity). (211 aa) |
| LTBP1 | Latent transforming growth factor beta binding protein 1. (1705 aa) |
| LAMA4 | Laminin subunit alpha 4. (1732 aa) |
| PRELP | Proline and arginine rich end leucine rich repeat protein. (380 aa) |
| SPARC | SPARC; Appears to regulate cell growth through interactions with the extracellular matrix and cytokines. Binds calcium and copper, several types of collagen, albumin, thrombospondin, PDGF and cell membranes. There are two calcium binding sites; an acidic domain that binds 5 to 8 Ca(2+) with a low affinity and an EF-hand loop that binds a Ca(2+) ion with a high affinity (By similarity); Belongs to the SPARC family. (315 aa) |
| COL1A2 | Collagen alpha-2(I) chain; Type I collagen is a member of group I collagen (fibrillar forming collagen); Belongs to the fibrillar collagen family. (1364 aa) |
| LAMC1 | Laminin subunit gamma 1. (998 aa) |
| COL4A1 | Collagen type IV alpha 1 chain. (1637 aa) |
| COL4A2 | Collagen type IV alpha 2 chain. (1564 aa) |
| HTRA1 | HtrA serine peptidase 1. (338 aa) |
| LAMB1 | Laminin subunit beta 1. (1736 aa) |
| COL5A2 | Collagen type V alpha 2 chain. (1502 aa) |
| MGP | Matrix Gla protein; Associates with the organic matrix of bone and cartilage. Thought to act as an inhibitor of bone formation. (103 aa) |
| HSD17B12 | Hydroxysteroid 17-beta dehydrogenase 12; Belongs to the short-chain dehydrogenases/reductases (SDR) family. (312 aa) |
| HSPG2 | Heparan sulfate proteoglycan 2. (4207 aa) |
| G1TVW1_RABIT | Fe2OG dioxygenase domain-containing protein. (273 aa) |
| TNC | Tenascin C. (2384 aa) |
| PXDN | Peroxidasin. (1411 aa) |
| THBS2 | Thrombospondin 2. (1170 aa) |

| | |
|---|---|
| ASPN | Asporin. (373 aa) |
| BGN | Biglycan; May be involved in collagen fiber assembly. (394 aa) |
| CRISPLD2 | Cysteine rich secretory protein LCCL domain containing 2. (471 aa) |
| FBLN5 | Fibulin 5. (453 aa) |
| COL5A1 | Fibrillar collagen NC1 domain-containing protein. (383 aa) |
| MMP2 | 72 kDa type IV collagenase; Ubiquitinous metalloproteinase that is involved in diverse functions such as remodeling of the vasculature, angiogenesis, tissue repair, tumor invasion, inflammation, and atherosclerotic plaque rupture. As well as degrading extracellular matrix proteins, can also act on several nonmatrix proteins such as big endothelial 1 and beta- type CGRP promoting vasoconstriction. Also cleaves KISS at a Gly-\|-Leu bond. Appears to have a role in myocardial cell death pathways. Contributes to myocardial oxidative stress by regulating the activity of GSK3beta. Cleaves GSK3 [...] (710 aa) |
| GPC6 | Glypican 6; Cell surface proteoglycan that bears heparan sulfate. Belongs to the glypican family. (555 aa) |
| ENSOCUP00000037812 | Uncharacterized protein. (1162 aa) |
| NID1 | Uncharacterized protein. (1000 aa) |
| LOX | Lysyl oxidase. (419 aa) |
| ENSOCUP00000044803 | Uncharacterized protein. (811 aa) |
| ENSOCUP00000047220 | Uncharacterized protein. (573 aa) |
| **Predicted functional proteins** | |
| COL3A1 | Collagen type III alpha 1 chain. |
| POSTN | Periostin. |
| DCN | Decorin; May affect the rate of fibrils formation; Belongs to the small leucine-rich proteoglycan (SLRP) family. SLRP class I subfamily. |
| COL1A1 | Collagen alpha-1(I) chain. |
| ENSOCUP00000040886 | Uncharacterized protein. |
| FN1 | Fibronectin; Fibronectins bind cell surfaces and various compounds including collagen, fibrin, heparin, DNA, and actin. Fibronectins are involved in cell adhesion, cell motility, opsonization, wound healing, and maintenance of cell shape. Involved in osteoblast compaction through the fibronectin fibrillogenesis cell-mediated matrix assembly process, essential for osteoblast mineralization. Participates in the regulation of type I collagen deposition by osteoblasts (By similarity). |
| DPT | Dermatopontin. |
| COL4A3 | Collagen type IV alpha 3 chain. |
| COL12A1 | Collagen alpha-1(XII) chain; Type XII collagen interacts with type I collagen-containing fibrils, the COL1 domain could be associated with the surface of the fibrils, and the COL2 and NC3 domains may be localized in the perifibrillar matrix; Belongs to the fibril-associated collagens with interrupted helices (FACIT) family. |
| COL15A1 | Collagen type XV alpha 1 chain. |

**Table S2.** List of cell-extracellular matrix adhesion proteins identified in acellular fibers using proteomics analysis, generated using the STRING database.

| **Detected proteins** | |
|---|---|
| EMILIN1 | Elastin microfibril interfacer 1. (982 aa) |
| LAMC1 | Laminin subunit gamma 1. (998 aa) |
| LAMB1 | Laminin subunit beta 1. (1736 aa) |
| TNC | Tenascin C. (2384 aa) |
| **Predicted functional proteins** | |
| ITGA9 | Integrin_alpha2 domain-containing protein; Belongs to the integrin alpha chain family. |
| ITGA10 | Integrin subunit alpha 10; Belongs to the integrin alpha chain family. |
| ITGA7 | Integrin subunit alpha 7; Belongs to the integrin alpha chain family. |
| ITGA1 | Integrin subunit alpha 1; Belongs to the integrin alpha chain family. |
| ITGA11 | Integrin subunit alpha 11; Belongs to the integrin alpha chain family. |
| ITGA4 | Integrin subunit alpha 4; Belongs to the integrin alpha chain family. |
| ITGA2 | Integrin subunit alpha 2; Belongs to the integrin alpha chain family. |
| ITGB8 | Integrin beta-8; Integrin alpha-V:beta-8 (ITGAV:ITGB8) is a receptor for fibronectin (By similarity). It recognizes the sequence R-G-D in its ligands (By similarity). Integrin alpha-V:beta-6 (ITGAV:ITGB6) mediates R-G-D-dependent release of transforming growth factor beta-1 (TGF-beta- 1) from regulatory Latency-associated peptide (LAP), thereby playing a key role in TGF-beta-1 activation on the surface of activated regulatory T-cells (Tregs) (By similarity). Required during vasculogenesis (By similarity). |
| ITGAV | Integrin subunit alpha V; Belongs to the integrin alpha chain family. |
| SV2C | Synaptic vesicle glycoprotein 2C. |

**Table S3.** Number of animals per conditions and time point.

| **Time point** | **Test** | **Ctrl** | **Sham** |
|---|---|---|---|
| Week 2 | 5 | 4 | 5 |
| Week 4 | 5 | 5 | 4 |
| Week 8 | 6 | 5 | 4 |

**Table S4.** List of primary and secondary antibodies used for immunohistochemistry.

| **Primary antibodies** | | |
|---|---|---|
| **Target** | **Dilution** | **Source** |
| Rabbit anti-Laminin | 1:500 | Abcam, ab11575 |
| Rabbit anti-Desmin | 1:300 | Abcam, ab32362 |
| Rabbit anti-Collagen I | 1:400 | Abcam, ab270993 |
| Rabbit anti-FSP1 | 1:200 | Sigma, ABF32 |
| Rabbit anti-CD31 | 1:200 | Abcam, ab182981 |
| Rabbit anti-CD68 | 1:100 | R&D Systems, MAB101141 |
| **Secondary antibodies** | | |
| **Target** | | **Source** |
| Peroxidase conjugated donkey anti mouse antibody | | Jackson ImmunoResearch, 715-035-150 |
| Peroxidase conjugated donkey anti mouse antibody | | Jackson ImmunoResearch, 715-035-152 |

**Figure S1.** Protein-protein interaction network analysis of extracellular matrix components visualized using K-means clustering (k=3) generated using the STRING database. Line thickness indicates the strength of data support for known associations.

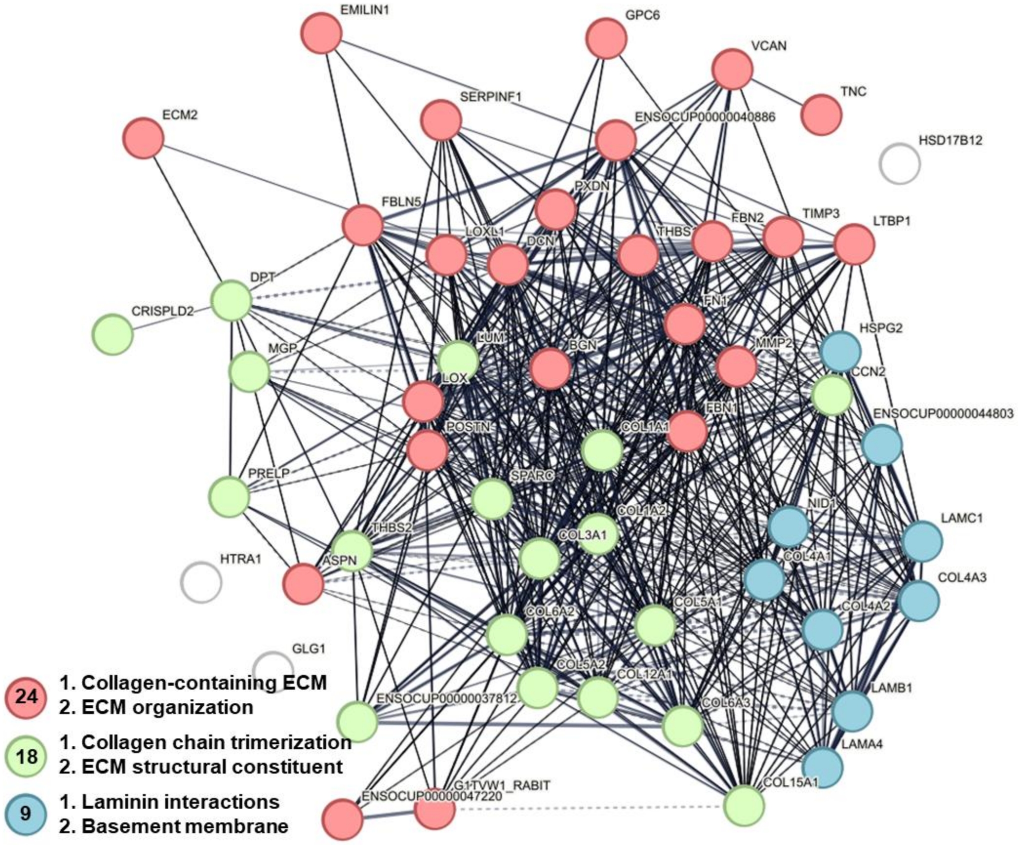


**Figure S2.** Protein-protein interaction network analysis of cell-ECM adhesion proteins visualized using K-means clustering (k=3) generated using the STRING database. Line thickness indicates the strength of data support for known associations.

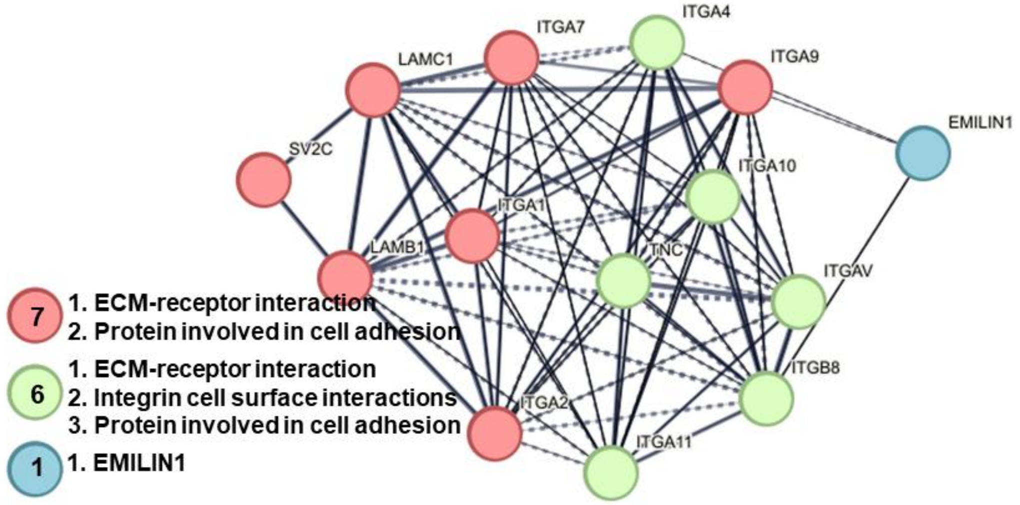

**Figure S3.** Detection of three zones in the muscle in QuPath software, actively remodeling treatment site, intact muscle and transition zone separating the first two. The transition zone could be muscle tissue being damaged due to inflammatory response to the damage or new partially regenerated muscle. As it couldn't be confidently assigned to either, only the actively remodeling site was assessed in the current study.

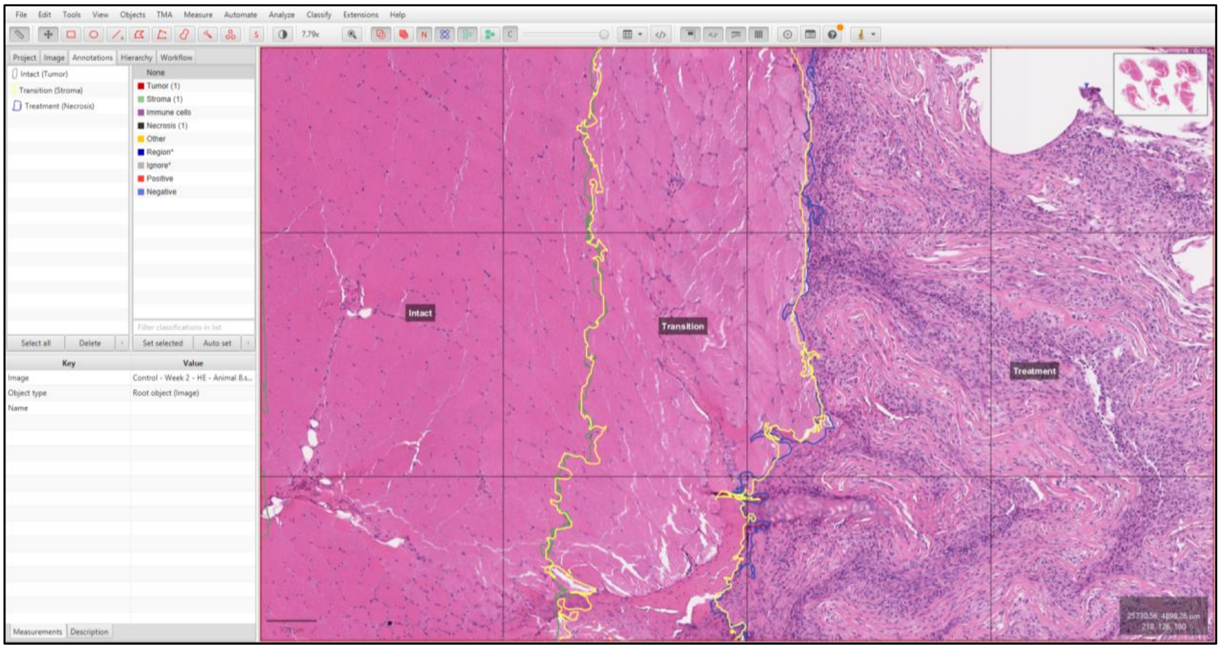


**Figure S4.** Force-frequency relationship of dorsiflexor muscles under different experimental conditions. Native tissue shows highest force production, with peak forces reaching ~1300 mN at maximal stimulation frequency (150 Hz). Error bars represent standard error of the mean (SEM). n is 6, 6, 5, and 4 for native (contralateral muscle), test, control, and sham groups, respectively.

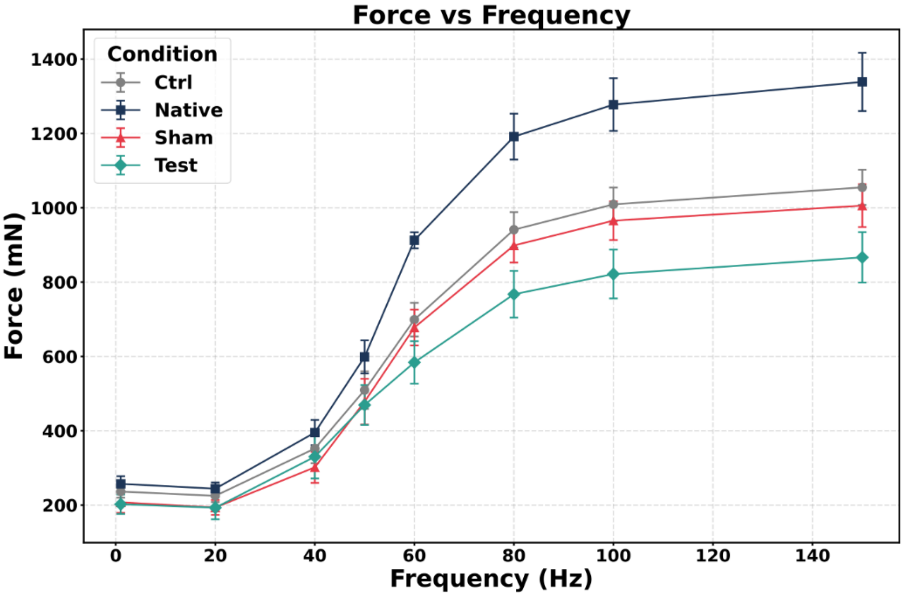

**Figure S5.** Comprehensive statistical analysis for histology image analysis. A mixed-effects model was employed followed by a pairwise t-tests with unequal variance assumption (Welch's t-test) between groups at different timepoint. P-values were adjusted for multiple comparisons using the Bonferroni correction method.

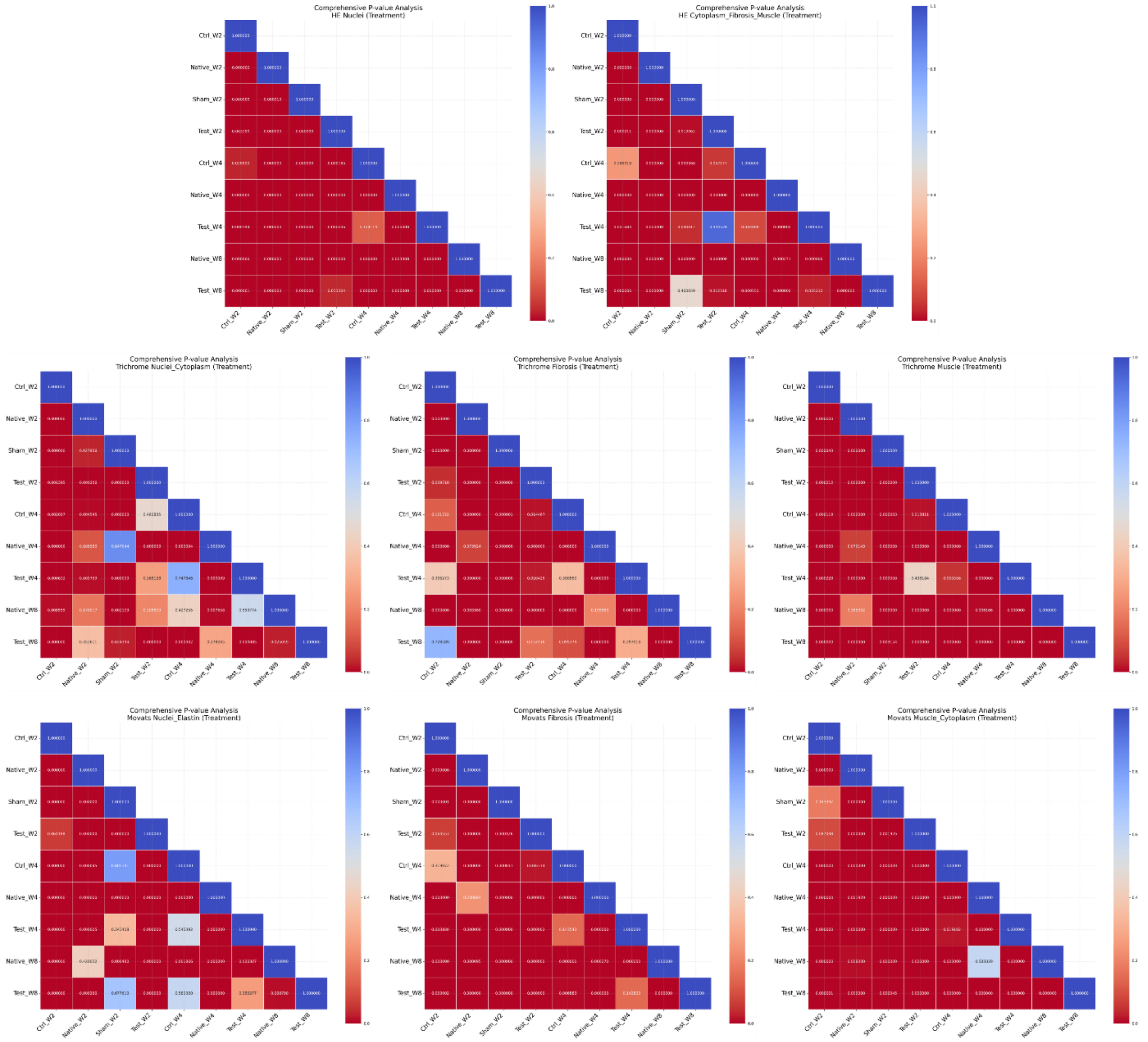

**Figure S6.** Comprehensive statistical analysis for IHC images using slide- and tile-level values. For tile-level data the same mixed-effects model was employed. Slide-level data (mean of all tile values for each slide) were used with a weighted two-sample t-tests with Bonferroni correction implemented to account for standard deviations calculated from tile level data.

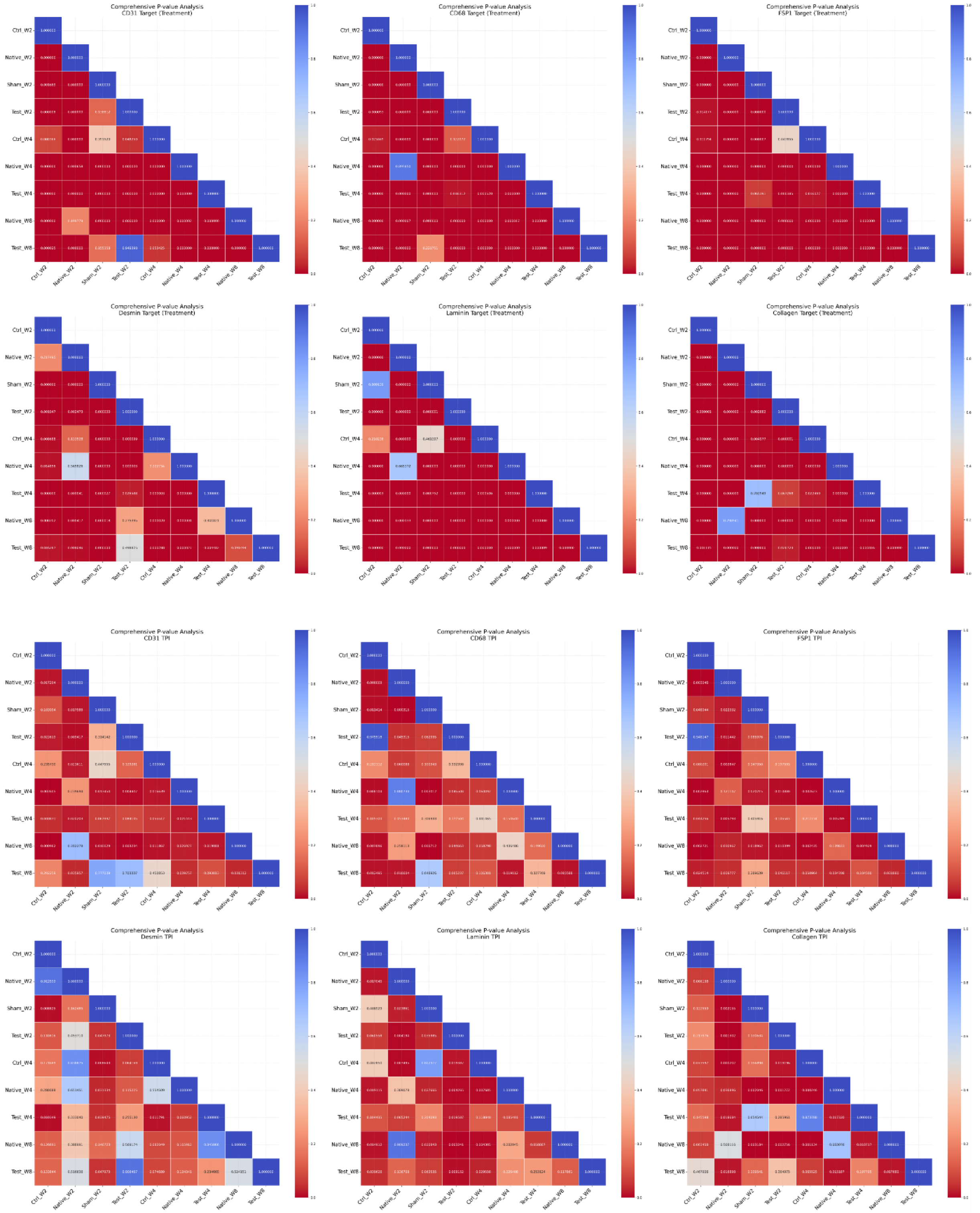
